# The *Wolbachia* WalE1 effector alters *Drosophila* endocytosis

**DOI:** 10.1101/2023.02.26.530160

**Authors:** MaryAnn Martin, Irene L.G. Newton

## Abstract

The most common intracellular bacterial infection is *Wolbachia pipientis*, a microbe that manipulates host reproduction and is used in control of insect vectors. Phenotypes induced by *Wolbachia* have been studied for decades and range from sperm-egg incompatibility to male killing. How *Wolbachia* alters host biology is less well understood. Previously, we characterized the first *Wolbachia* effector – WalE1, which encodes a synuclein domain at the N terminus. Purified WalE1 sediments with and bundles actin and when heterologously expressed in flies, increases *Wolbachia* titer in the developing oocyte. In this work, we first identify the native expression WalE1 by *Wolbachia* infecting both fly cells and whole animals. WalE1 appears as aggregates, separate from *Wolbachia* cells. We next show that WalE1 co-immunoprecipitates with the host protein Past1 and that WalE1 manipulates host endocytosis. Yeast expressing WalE1 show deficiency in uptake of FM4-64 dye, and flies harboring mutations in *Past1* or overexpressing WalE1 are sensitive to AgNO_3_, a hallmark of endocytosis defects. Finally, we also show that *Past1* null flies harbor more *Wolbachia* overall and in late egg chambers. Our results identify interactions between a *Wolbachia* secreted effector and a host protein and point to yet another important host cell process impinged upon by *Wolbachia*.

## Introduction

*Wolbachia pipientis* is an obligate intracellular microbe and arguably the most successful infection on our planet, colonizing 40-60% of insect species, as well as other arthropods and filarial nematodes (1, 2). *Wolbachia* are alpha-proteobacteria, part of the anciently intracellular *Anaplasmataceae*, and related to the important human pathogens *Anaplasma, Rickettsia* and *Ehrlichia* (3). However, *Wolbachia* do not infect mammals, but instead are well known for their reproductive manipulations of insect populations, inducing phenotypes such as male-killing, feminization, or sperm-egg incompatibility (4). In the last decade, *Wolbachia* have also been shown to provide a benefit to insects, where the infection can inhibit RNA virus replication within the host (5), a phenomenon known as pathogen blocking. Because insects are vectors for disease, and *Wolbachia* alter the ability of these vectors to harbor important human pathogens, *Wolbachia* are being used to control the spread of diseases such as dengue.

Like all intracellular bacteria, *Wolbachia* need to manipulate the host cell to invade and persist. Many microbes accomplish this via secretion systems, nanomachines that enable the microbes to directly transfer proteins from the bacterium into the cytosol of host cells. Based on analyses of the genomes of all sequenced strains, *Wolbachia* symbionts encode a functional type IV secretion system (T4SS) (6-8), which is expressed by *Wolbachia* within its native host (7, 9). However, only a few proteins secreted by *Wolbachia* have been identified and characterized. These proteins, referred to as effectors, often act to manipulate or usurp host cell processes to promote bacterial infection (10, 11). These modes include (but are not limited to) attacking the host cell surface to form pores, inactivating host cytosol machinery to collapse the cytoskeleton, or entering the nucleus to manipulate host gene regulation (12). At each stage of attack, the bacterial effectors often interact directly and specifically with host proteins to perturb a biological process that enables pathogen entry into or defense from the host cell (13-19). Understanding how bacterial effectors function, therefore, has taught scientists not only how pathogens cause disease, but also fundamental cell biological mechanisms work in healthy tissue (20). While effectors are bacterial in origin, they act within eukaryotic cells and hence often encode domains that share structural, functional, and sequence similarity with eukaryotic proteins (10, 11, 21, 22).

The first characterized *Wolbachia* effector, WalE1, is an actin bundling protein that increases *Wolbachia* titer in the next generation when over-expressed in transgenic flies (8). The WalE1 protein contains an N-terminal synuclein domain (8) which may mediate some interactions with host proteins and pathways. WalE1 is upregulated by *Wolbachia* during host pupation and purified WalE1 protein co-sediments with filamentous actin and increases actin bundling *in vitro* and *in vivo*. As the actin cytoskeleton is important for *Wolbachia’s* maternal transmission (23), and for its internalization by host cells (24), the WalE1 effector likely plays an important role in *Wolbachia’s* biology.

In this study, we sought to characterize the patterns of expression of native WalE1, its host targets (beyond actin) and identify specific host pathways influenced by the effector. We used antibody made to the full-length protein to visualize the effector during *Wolbachia* infection of *Drosophila* cells and ovaries. The native protein appears as large aggregates in host cells and early egg chambers. We use co-immunoprecipitation and mass spectrometry to determine host proteins with which WalE1 interacts and identify Past1 as a target of WalE1. Past1 was previously shown to influence endocytosis (25); Garland cells from homozygous *Past1* mutant larvae were defective in their ability to endocytose fluorescently labeled avidin. We therefore characterized endocytosis defects upon both WalE1 expression in yeast and flies and Past1 abrogation in flies. Finally, we show an interaction between *Wolbachia* titers and Past1 titers in whole animals, where *Wolbachia* titer increases in Past1 mutant flies, suggesting that *Wolbachia* targets the protein to alter its function. Our results shed light on the molecular mechanisms used by a ubiquitous symbiont to alter host biology.

## Results

### Native WalE1 localizes to aggregates in the host cell cytosol

We had previously shown phenotypes for overexpression of RFP-tagged WalE1 in whole flies (8). However, this experiment could have been affected by the use of tags and non-native expression; tags can alter protein localization and we were necessarily studying an artificial system and not native expression and secretion of the effector by *Wolbachia*. We therefore generated an antibody to full length WalE1 and used it to probe cells and flies infected with *Wolbachia*. The α-WalE1 antibody does not stain *Drosophila* JW18 cells cleared of their *Wolbachia* infection with tetracycline. However, in JW18 cells infected with strain *w*Mel, α-WalE1 localizes to large aggregates that do not colocalize with *Wolbachia* (stained with DAPI, Figure 1). In ovaries, we observe similar aggregates of WalE1 in *Wolbachia-*infected flies only. These aggregates appear most numerous in early stages of oogenesis (in the germarium and stages 2-4), when *Wolbachia* is most numerous (23), and seem to disappear in later egg chambers (Figure 2).

**Figure 1.**
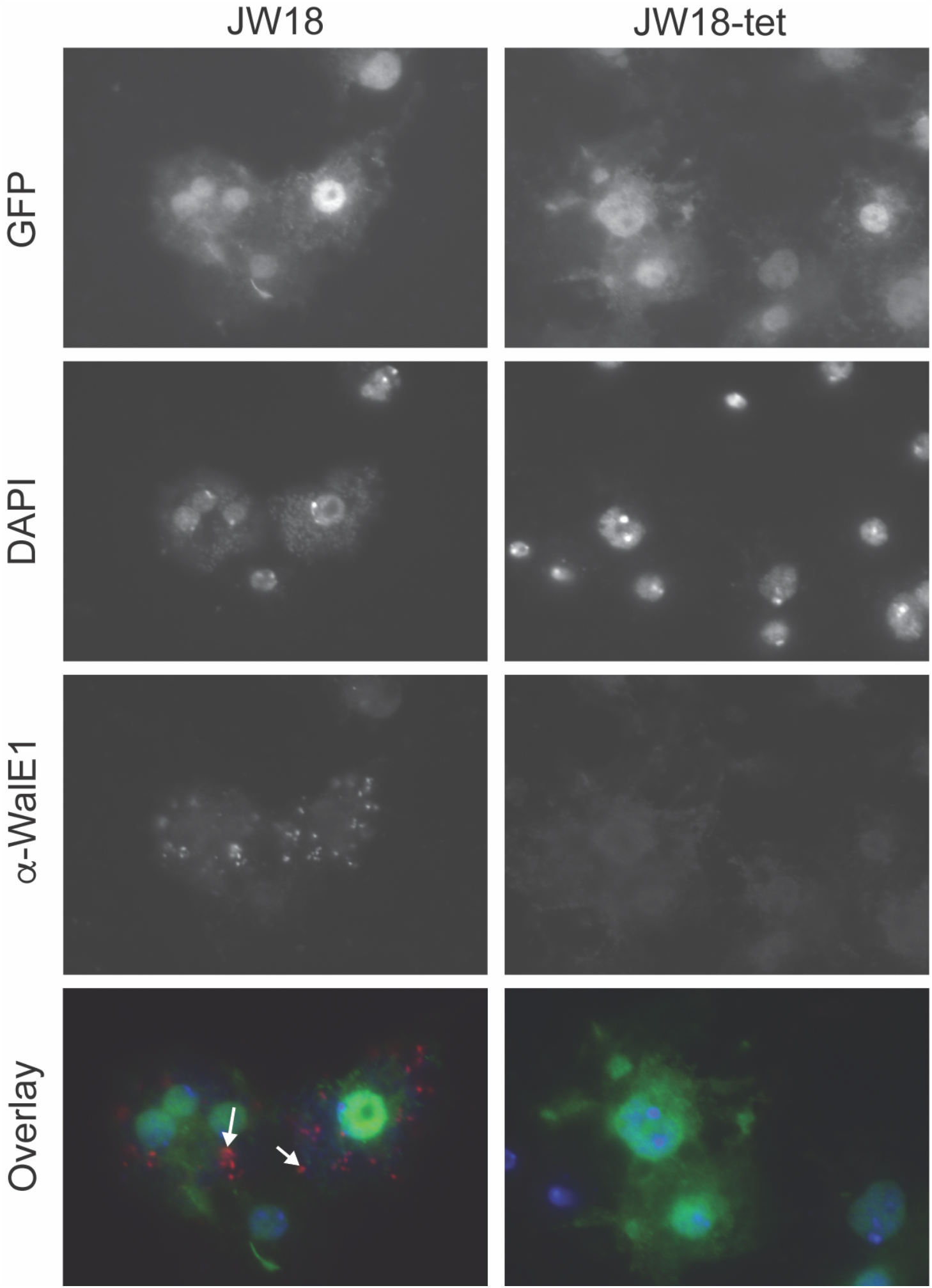
WalE1 native expression in *Wolbachia-*infected JW18 cells. Antibody made against full-length WalE1 was used to probe *Wolbachia-*infected (JW18) and uninfected (JW18-tet) cells. Native WalE1 expression is only seen in JW18 cells, where the protein localizes in large aggregates, outside of *Wolbachia* cells (stained with DAPI). Arrowheads point to WalE1 aggregates in overlay.

**Figure 2.**
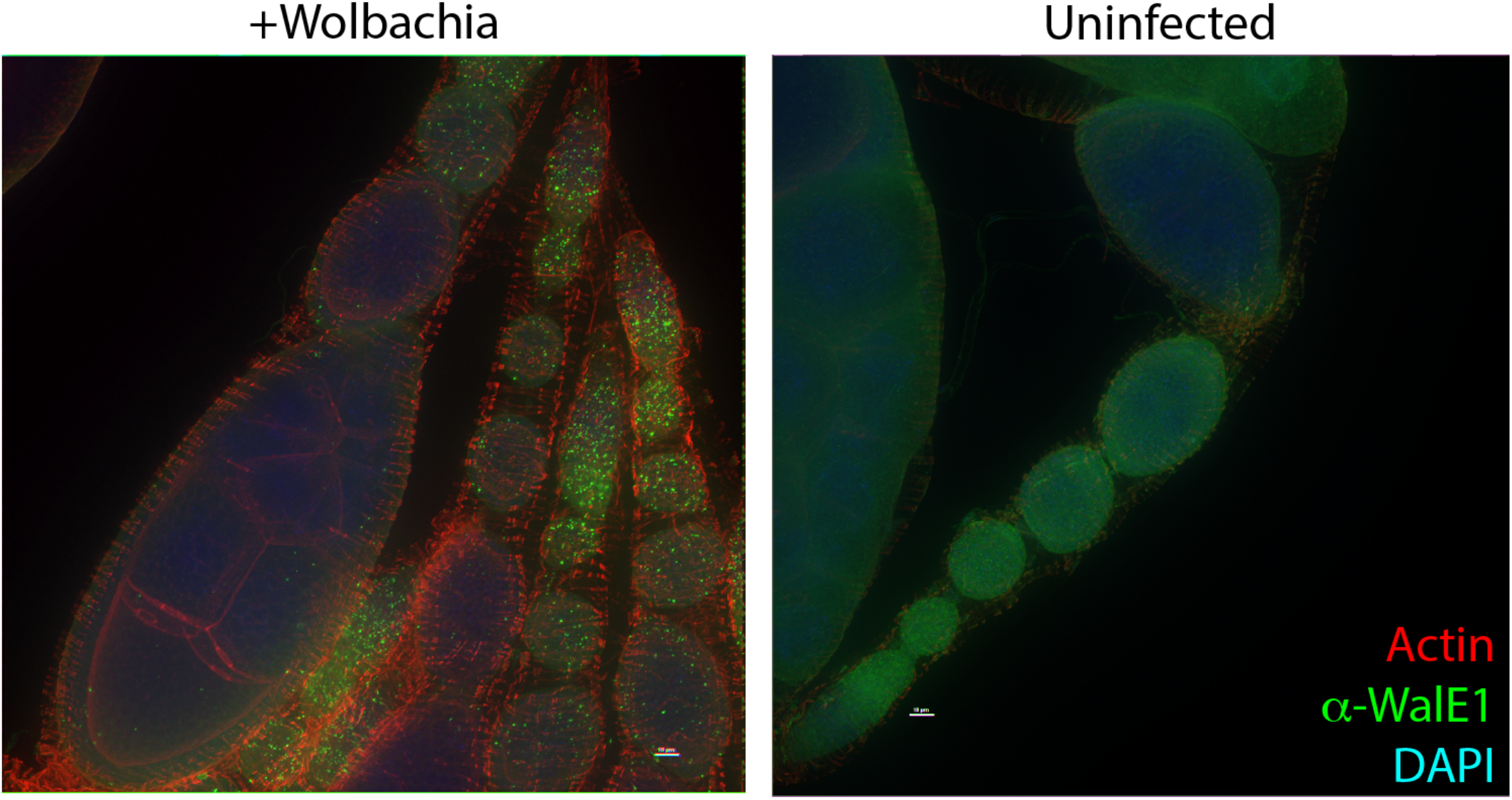
WalE1 native expression in *Wolbachia-*infected fly ovarioles. Antibody made against full-length WalE1 was used to probe *Wolbachia-*infected and uninfected ovaries. Native WalE1 expression is seen as green puncta in *Wolbachia*-infected germaria through stage 9 egg chambers.

### WalE1 interacts with host Past1 and the endocytosis pathway

Because native WalE1 localized to aggregates in the cell, we sought to identify what host proteins may be localized to those aggregates, interacting with WalE1. Towards that end, we performed a co-immunoprecipitation experiment using α-WalE1 antibody. Infected *Drosophila* JW18 cells were lysed and incubated with α-WalE1 antibody-coated magnetic beads. As controls we included beads alone and uninfected JW18 cells were also subjected to the same immunoprecipitation. After washes on a magnetic column, bound proteins were eluted and subjected to mass spectrometry. We identified one protein, significantly enriched in our pull down, which came down in the JW18 lysate but was absent from the JW18-tet lysate: Putative Achaete Scute Target 1 (Past1) (Table 1).

**Table 1.** Co-immunoprecipitation of WalE1 using α-WalE1 antibody pulls down host Past1 protein from *w*Mel infected cell line JW18, but not from uninfected cells.

| Protein Name | JW18 |  |  | JW18-TET |  |  |
| --- | --- | --- | --- | --- | --- | --- |
|  | Num Unique | Peptide Count | % Cov | Num Unique | Peptide Count | % Cov |
| Zipper, isoform D | 108 | 279 | 49.9 | 109 | 246 | 52.4 |
| RE59368p (Past1) | 17 | 21 | 35 |  |  |  |
| Short stop/Kakapo long isoform | 17 | 18 | 4.6 | 4 | 4 | 1 |
| Karst, isoform B | 18 | 18 | 5.3 | 6 | 6 | 1.8 |
| AP-3 complex subunit beta | 21 | 24 | 21.6 | 15 | 16 | 14.9 |
| Poly(U)-binding-splicing factor half pint | 9 | 12 | 18.5 | 6 | 6 | 14.8 |

Past1 is an EHD (Eps15 Homology Domain) ortholog in *Drosophila* and known to contribute to endocytosis (25). Eps15 Homology Domain containing proteins in other models are involved in the recycling of proteins and lipids to the plasma membrane (26-30). Indeed, Past1 has previously been shown to play a role in endocytosis; Garland cells from *Past1*^*110-1*^ homozygous null flies do not endocytose fluorescently labeled avidin to the same extent as wildtype (25). As WalE1 interacts with Past1, we therefore wondered if WalE1 expression might also impact endocytosis. We used FM4-64 labeling of yeast to measure extent of endocytosis in a pulse chase experiment. Yeast expressing WalE1 internalize less FM4-64 than control yeast carrying vector alone (Figure 3, χ2= 24.706, df = 1, p = 6.676e-07). This result suggests that WalE1 expression may impact endocytosis, likely through interaction with Past1. We next moved our analyses to *Drosophila melanogaster* Past1.

**Figure 3.**
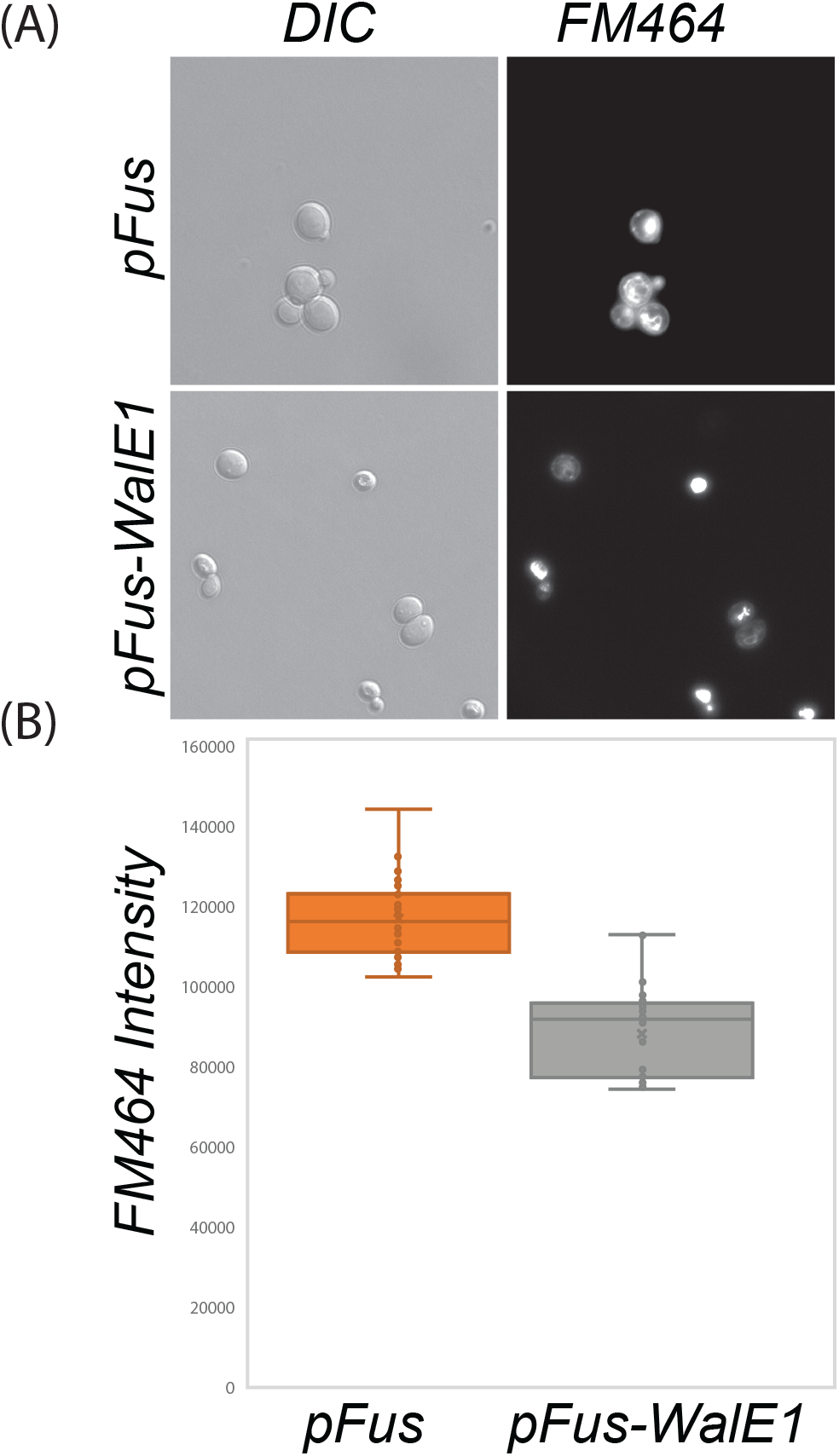
Yeast expressing WalEl internalize less FM464 dye than control yeast carrying vector alone. Yeast carrying control vector or expressing *WalE1* were exposed to FM464 membrane dye for a pulse and internalization of dye was quantified after 60 minutes based on fluorescence intensity (N > 40 for each; χ^2^ = 24.706, df = 1, p = 6.676e-07).

### Past1 mutant flies do not suffer fertility or viability defects

Past1 mutants were previously characterized to suffer from fertility defects, exhibit sensitivity to temperature, and die early (25). We began by acquiring the most commonly used alleles (*Past1*^*60-4*^ and *Past1*^*110-1*^); both of these alleles were made by imprecise excision of a p-element insertion, which result in *Past1* nulls (25). Our first western blot experiments confirmed that these mutants are indeed nulls for Past1 (Figure S1). We next focused on using deficiency stocks (Df), containing chromosomal ablations, covering the Past1-containing region, to confirm phenotypes previously published for these alleles (25). The Df used in the previous study was not available to us (Df(3R)Kar-Sz37 87C5-87D14), so we used two deficiencies from the Bloomington Drosophila Stock Center, both of which covered *Past1* entirely, and the larger of which (Df(3R)BSC486) led to greater levels of lethality. The smaller deficiency (Df(3R)Kar-Sz29) allowed for high levels of viability in animals lacking *Past1*, and in our extensive study of ovaries from these flies to study *Wolbachia* titer effects, oogenesis defects were never observed (Figure 4). These results suggest that something else in the background of the *Past1*^*60-4*^ and *Past1*^*110-1*^ flies contributes to the viability and fertility defects previously observed. Importantly, prior work did not specify if the allele used (*Past1*^*110-1*^) was in a *white-* background. The P-element used (EY01852) carries mini-white^+^, and white mutants impair several biological functions – from mobility to lifespan to stress tolerance (31). Therefore, it is entirely possible that the viability and fertility defects previously published were confounded by the background of these mutations.

**Figure 4.**
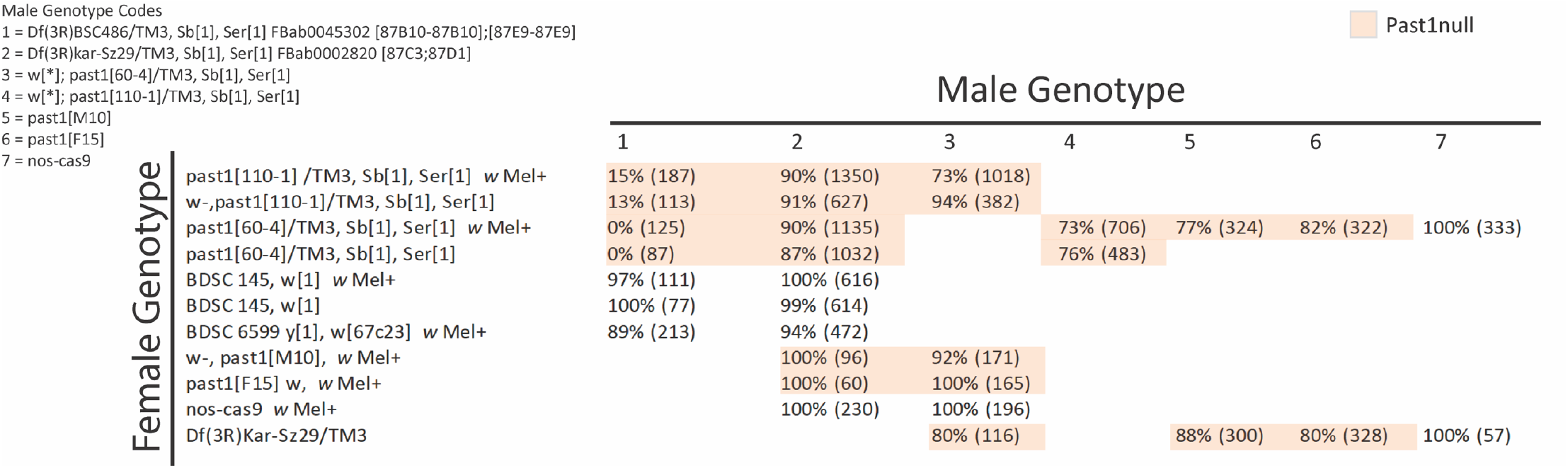
Past1 null mutants are viable. Male genotype for each cross is shown at top right, and females on the lefthand side. Total number of flies counted is noted in parentheses and viability is calculated as a percentage. Severely reduced viability is observed with the large deficiency Df(3R)BSC486, which covers adjacent genes, but other combinations of hemizygous *past1* null alleles lead to full or higher percentages of viable offspring. Past1 nulls shown in light orange.

We sought to genetically isolate any effects that may be related to *Past1* in our assays and therefore generated new CRISPR-Cas9 based mutants in Past1 (*Past1*^*M10*^ and *Past1*^*F15*^). These mutants generate an identical two basepair deletion that generates a frameshift and eventual stop codon after 83 amino acids, resulting in production of no stable protein (Figure S1). We used transheterozygotes of the previously generated allele *Past1*^*60-4*^ and our two CRISPR generated null alleles (*Past1*^*M10*^ and *Past1*^*F15*^) in these experiments. Our genetic controls were progenitor flies from the Cas9 process, that are wild-type for Past1.

### Past1 mutant flies exhibit endocytosis defects

Previously, endocytosis defects were observed for *past1* mutant flies (25). In that prior study, the authors had used *Drosophila* Garland cells to evaluate endocytosis in the fly as they are the equivalent of a kidney, filtering the fly haemolymph. One straightforward way to test for fly Garland cell function is to expose flies to silver nitrate (AgNO_3_); if the fly harbors mutations that alter function of the Garland cells, they will be more sensitive to AgNO_3_ toxicity (32). We therefore subjected larvae expressing *walE1* and larvae carrying mutations in *Past1* to AgNO_3_ throughout all of development and monitored viability over time, counting the number of pupae and adults derived from these conditions. Control flies did not show any phenotype when reared in the presence of AgNO_3_ and indeed, no statistical differences were observed across all fly backgrounds reared in normal food without AgNO_3_. However, flies expressing the *Wolbachia* effector were exquisitely sensitive to AgNO_3_, to the same extent as those carrying *Past1* null mutations (Figure 5). While 90% of control flies survive AgNO_3,_ WalE1 overexpression resulted in 5% survival on average (Figure 5). This result confirms prior observations regarding the effect of Past1 on endocytosis (25) and supports a role for WalE1 in modifying endocytosis as well. Importantly, all flies used in this experiment did not have the *white* mutation nor balancer chromosomes, to remove confounding background effects (Supplementary Table 2).

**Figure 5.**
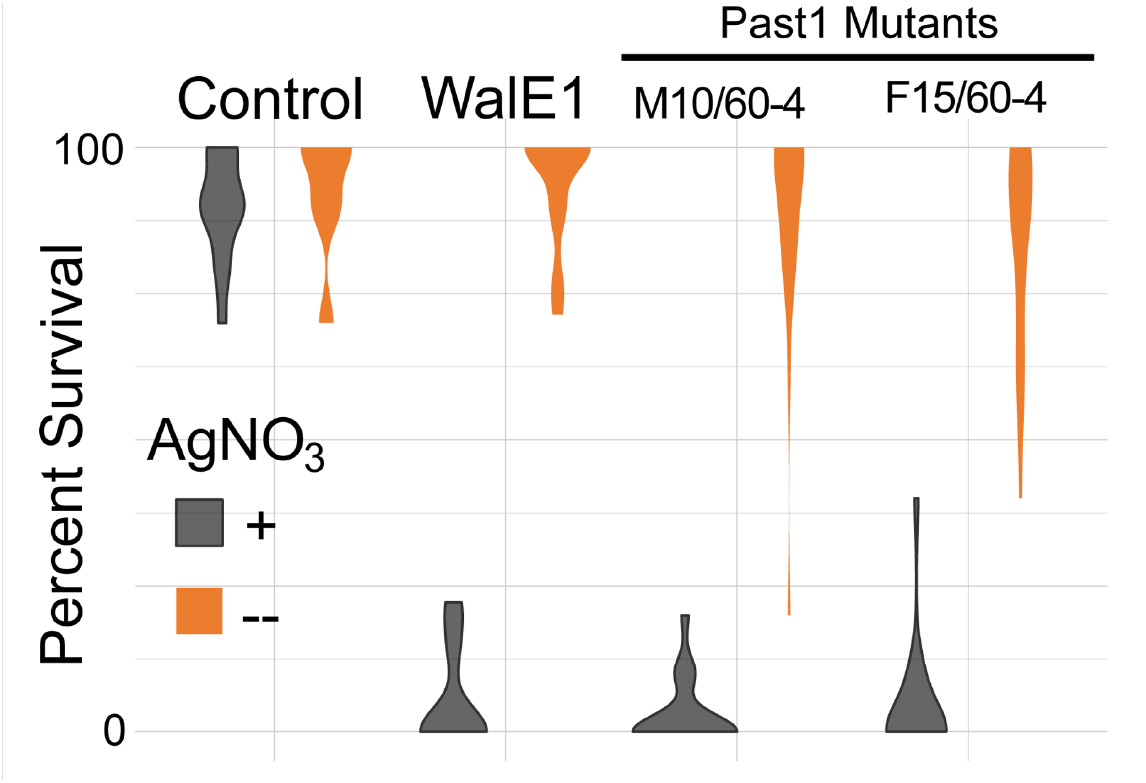
Survival of adult flies subjected to silver nitrate exposure during development. Flies expressing WalE1 or carrying mutations in Past1 are sensitive to AgNO_3_ (gray) exposure compared to survival on control food (orange). Control flies show no lethal phenotype. N > 100, across 5 separate vials for each condition.

### Wolbachia attain higher titer in Past1 null flies

Because we identified Past1 as a potential target of WalE1, and because both proteins are involved in modifying endocytosis in *Drosophila*, we wondered if there would be an interaction between *Past1* null mutants and *Wolbachia* titer. We reasoned that if *Wolbachia* was using WalE1 to modify Past1, reduction of the dosage of *Past1* might influence *Wolbachia* biology. Therefore, we began by examining *Wolbachia* titer in flies with different copy numbers of *Past1*. We used western blot targeting the *Wolbachia* surface protein and observed a clear and statistically significant effect of *Past1* dosage on *Wolbachia* titer (ANOVA; df = 2, F = 40.611, p < 0.001). Flies that are null for *Past1* have the highest *Wolbachia* titer (t = -7.675, df = 13, p < 0.001) followed by flies with a half dose of *Past1*, although the titer in these flies is not significantly different from wild-type (t = -2.632, df = 10, p = 0.457) (Figure 6).

**Figure 6.**
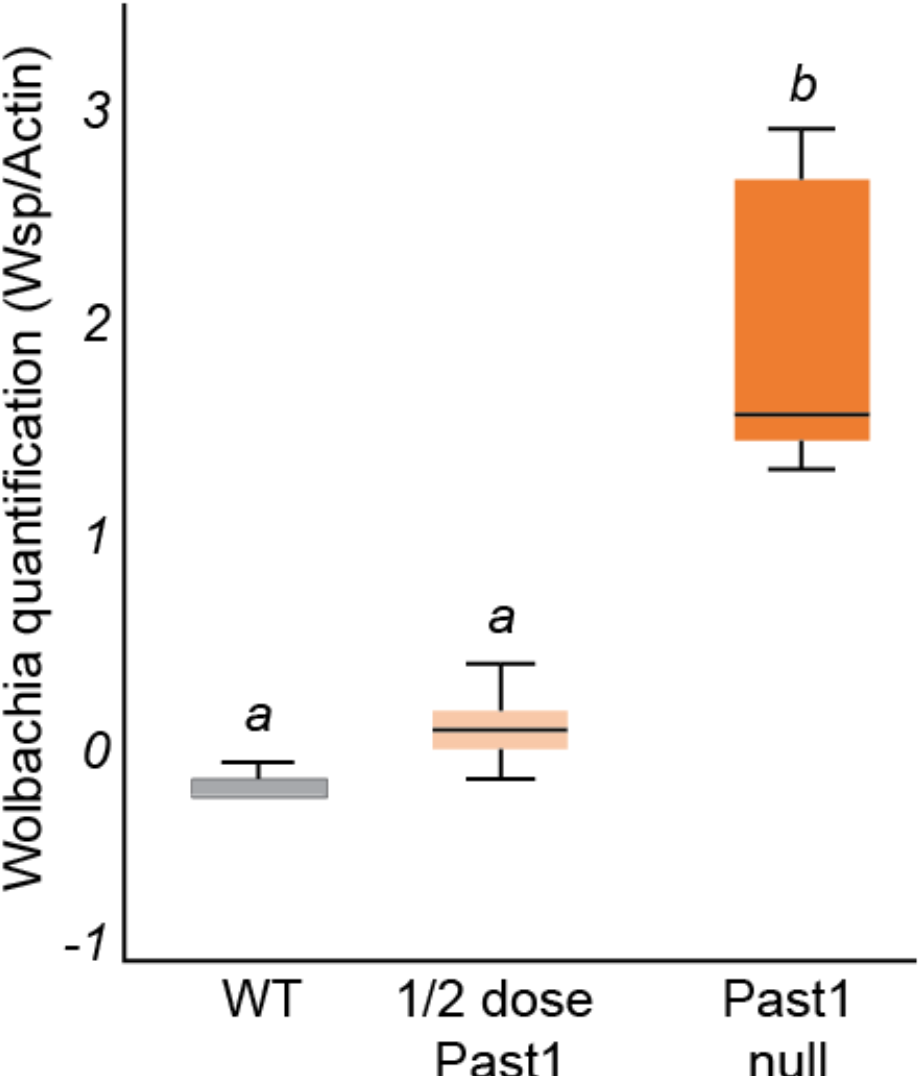
*Wolbachia* titer increases in flies in a Past1 dose-dependent manner. Entire flies were used in a western-blot targeting *Wolbachia’s* surface protein (WSP) band intensity was measured relative to α-Actin band intensity as a loading control. *Wolbachia* titers increase in whole flies and are highest in the Past1 null flies (*Past1^60 4^/Df(3R*)Kar-Sz29), intermediate in titer in ½ dose Past1 flies (*Past1*^*60-4*^ *or Df(3R)Kar-Sz29*/*Past1*^*+*^ on balancer) flies, and lowest titer in flies with two copies of *Past1*^*+*^ (WT, wild-type flies) (ANOVA: df = 2, F = 40.611, p < 0.001).

We next wondered if this whole-animal increase in *Wolbachia* load would be observed in the ovary tissue as well, the tissue best studied for *Wolbachia* titer and localization. We dissected ovaries from *Past1*^*60-4*^ /Df(3R)Kar-Sz29 flies and sibling controls with a half dose of *Past1. Wolbachia* titer was quantified by *Wolbachia-*specific antibody staining (α-FtsZ) in ovarioles as indicated in the methods. We noticed that *Wolbachia* staining was most intense in the germarium for both backgrounds (as expected) but as oogenesis progressed, flies null in *Past1* harbored more *Wolbachia* in older egg chambers (Figure 7, GLM χ2 = 10.477; df = 3; p = 0.001). The most differentiation in *Wolbachia* titer between these backgrounds was observed for stages 7-8 (Figure 7; GLM χ2 = 39.449; df = 3; p < 0.001). This result is reminiscent of the increase in *Wolbachia* titer observed upon over-expression of WalE1 in our prior work (8).

**Figure 7.**
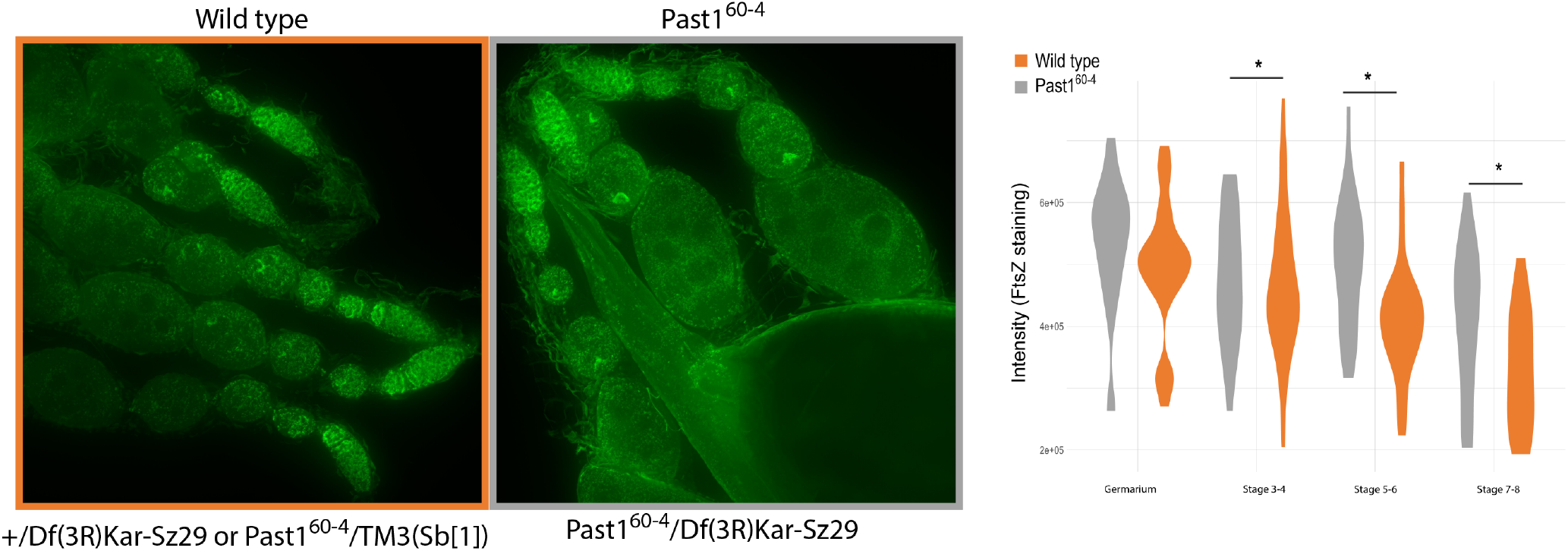
*Wolbachia* titer within the ovary of *Past1* mutant flies increases during oogenesis. *Past1*^*60-4*^ /Df(3R)Kar-Sz29 flies compared to their wild type counterparts (sibling controls with a null mutant copy of *Past1* and a balancer with *Past1*^*+*^). Wolbachia titer quantified by α-FtsZ staining in ovarioles (GLM χ^2^ = 10.477; df = 3; p = 0.001).

## Discussion

*Wolbachia* is a ubiquitous intracellular symbiont, infamous for reproductive manipulations induced in insects, such as cytoplasmic incompatibility and male killing (33). However, all the dramatic effects induced by *Wolbachia* require the bacterium to infect and persist in the insect host. Although a few proteins facilitating *Wolbachia’s* reproductive manipulations have been identified (34), we know even less about *Wolbachia’s* basic biology and how it maintains its infection. The first characterized *Wolbachia* effector was WalE1, an actin-bundling protein that increased *Wolbachia* titer in the oocyte upon overexpression (8). Purified WalE1 sediments with actin, and when heterologously expressed in flies and yeast, co-localizes with actin filaments (8). Here, we explored the native expression of WalE1 during a *Wolbachia* infection in *Drosophila* and find that it forms aggregates within both cell lines and oocytes. We characterized host proteins that may interact with WalE1 using co-immunoprecipitation and mass spectrometry and identified Past1 as a putative WalE1 target. We confirmed that *Drosophila* Past1 plays a role in endocytosis, and showed that WalE1 plays a similar role in both yeast and flies. Finally, we show that *Wolbachia* titer increases in egg chambers within Past1 null flies. This result echoes our previously published work showing that overexpression of WalE1 increased *Wolbachia* titers in the developing oocyte (8).

Here we show for the first time that a *Wolbachia*-secreted effector impacts host endocytosis. Many bacterial pathogens deploy effectors that alter host endocytosis and vesicle trafficking, as these cellular processes are important for signaling, interactions with the extracellular environment, and metabolism. By altering host endocytosis and vesicle trafficking, pathogens can avoid phago-lysozome fusion, as *Mycobacterium tuberculosis* does using EsxH (35), or even induce uptake of the microbe, as is the case for *Helicobacter pylori’s* CagA (36). The *Legionella pneumophila* effector AnkX, which encodes a FIC domain that transfers a phosphocholine to Rab1b, interferes with host endocytic recycling and localizes to tubular membrane compartments (37). Similarly, *Shigella* alters the composition of lipids in the vacuolar membrane – especially phosphoinositide composition – altering the stability of the *Shigella*-containing vacuole (38). We do not yet know exactly how *Wolbachia’s* WalE1 might modify Past1, or if the interaction is direct. However, we know that expression of WalE1 alters host endocytosis and that the *Wolbachia* protein pulls down host Past1, which is also implicated in endocytic recycling. What specific downstream cell biology is impacted by WalE1-Past1 interaction – and how this affects *Wolbachia* – is part of future work.

Although we have characterized native WalE1 and a putative target, many questions remain. As EHD proteins form large oligomers on membranes, does WalE1 co-localize with Past1? Unfortunately, our Past1 antibody, useful for western blot, did not function in immunohistochemistry, making that question more difficult to answer at the moment. EDH proteins like Past1 are involved in trafficking but we don’t know if WalE1 is targeting Past1 to impact specific cargo. What we’ve shown is that Past1 null flies harbor higher *Wolbachia* titers, suggesting that WalE1 antagonizes Past1 to promote *Wolbachia* infection. We hypothesize that by abrogating Past1 function, WalE1 alters the endocytic recycling of membrane-associated receptors or proteins important for *Wolbachia* infection. The WalE1 effector is conserved across a wide swathe of the *Wolbachia* genus (8), and is found within both insect-associated *Wolbachia* strains and those that infect nematodes. It would be interesting to know if these other homologs function similarly in their respective hosts – bundling actin and affecting host endocytosis – or if their functions have diverged. Also, Past1 is the only EHD protein in the *Drosophila* genome. When *Wolbachia* infects other insects, does WalE1 target their Past1 homolog(s)? Our work begins to shed some light on how *Wolbachia* impinges upon host biology and cell biological targets of *Wolbachia* effectors.

## Materials and Methods

### Yeast WalE1 expression and endocytosis assay

Previously published yeast strains and constructs were used for this assay (8). Briefly, yeast strain S288C (BY4741 **MATa**) was transformed with sequence-verified pFus-WalE1 expression vectors generated above using the PEG/Lithium acetate method (39). Yeast transformants were inoculated into selective synthetic media with 2% (w/v) glucose. These cultures were grown overnight to saturation (at 30°C) before back dilution to an OD_600_ of 0.1 into media containing 2% galactose for induction of the construct for 6 hours. The cultures were harvested and kept at 4°C while FM4-64 (ThermoFisher) was added 1:50 to the cells for 10 minutes. Yeast cells were then washed and mounted on agar pads, placed at 30°C, and imaged live for a 2-hour time course with a 60x oil objective. FM464 staining intensity per image was determined using Nikon NIS Elements software.

### Western Blots

Proteins were separated on 4-20% Tris-Glycine NB precast minigels (NuSep) and transferred to PVDF membrane in Tris-Glycine transfer buffer with 15% methanol at 40v on ice for 3 hours. The membranes were blocked for 5 minutes in SuperBlock™ (TBS) Blocking Buffer (ThermoFisher Scientific), followed by incubation with antibodies diluted in SuperBlock™ (TBS) Blocking Buffer (for 1 hour at room temperature or overnight at 4°C) according to standard protocols. PageRuler Prestained Protein Ladder (ThermoFisher Scientific) was used as a molecular mass marker. Antibodies utilized include mouse anti-actin at 1:10,000 (Seven Hills Bioreagents, LMAB-C4); rabbit anti-Past1 at 1:1000 (from Mia Horowitz at Tel Aviv University (25)) and mouse anti-Wsp at 1:10,000 (BEI Resources, Inc. NR-31029). F(ab’)2-Goat anti-Rabbit IgG (H+L) (A24531) and goat anti-mouse IgG (G-21040) secondary antibodies conjugated to horseradish peroxidase (HRP) were used at 1:5,000 (ThermoFisher Scientific Invitrogen). SuperSignal West Pico Chemiluminescent Substrate was used to detect HRP on immunoblots per manufacturer instructions (ThermoFisher Scientific). Blots were re-probed after stripping (100mM Glycine, 0.15 ND-40, 1% SDS, pH 2) for 1 hour at room temperature, then overnight or up to three days at 4°C. The rabbit Past1 anti-sera also detects a protein band of ∼ 50 KD in *w*Mel-containing lysates that is not observed when the sera are preabsorbed with strips of PVDF membrane containing total *E. coli* lysate.

### Drosophila *cell and ovary immunochemistry and microscopy*

The *Drosophila melanogaster* JW18 cell line, naturally infected with *Wolbachia* strain *w*Mel, was used to visualize WalE1 in an infection (40). JW18-TET cells were passaged with 10 ug/ul tetracycline added to the culture medium and were used as an uninfected control cell line. Confluent monolayers were harvested from 25 cm_2_ non-vented tissue culture flasks, counted using a disposable hemocytometer (Fisher Scientific), and overlaid as 100 ul of 2 × 10_6_ or 200 ul of 1 × 10_6_ cells onto Concanavalin-A coated No. 1.5 coverslips (ConA 0.5 mg/ml applied and dried on to sterile, acid-washed coverslips and leaving a 2 mm ConA-free border) (41, 42). Cells were allowed to settle, attach, and spread out overnight before cells were fixed in 4% paraformaldehyde in PBS for 20 minutes, followed by four 1 mL washes of 1X PBST (0.2% Tween-20 added to 1X PBS). Coverslips were then exposed to blocking solution (PBST 0.2% Tween-20 and 0.5% BSA) for 15 minutes at room temperature before transfer to primary antibody (at concentrations indicated below) overnight in blocking solution at 4°C. In the morning, coverslips were washed in PBST and then allowed to incubate for 1-2 hours at room temperature in the dark in 100 microliters of secondary antibodies diluted in the blocking solution. Coverslips were washed for 5 minutes 3-4 times with one mL PBST, followed by a final dip in a 500 mL beaker of distilled water. Excess moisture was wicked off the edge of the coverslip with a tissue followed by mounting in 10 microliters of Prolong Gold Antifade Reagent with DAPI (Invitrogen) per coverslip on glass slides.

For immunolocalization of proteins in developing egg chambers, ovaries from mated four-day old females were dissected in cold PBS before transfer to 1.5 mL microcentrifuge tubes with 100ul Devitt’s solution (5.3% paraformaldehyde, 0.5% NP-40, 1X PBS) and 600 ul heptane (43). The ovary tubes were vigorously shaken by hand for 30 seconds to create an emulsion, before rocking for 10 minutes at room temperature. After three rinses of ovaries, gently pelleted by 5-second spins in a mini microcentrifuge (6000 rpm, 2,000g), in PBST (0.2% Tween-20 added to 1X PBS), the ovaries were soaked in blocking solution (PBST 0.2% Tween-20 and 0.5% BSA) for 15 minutes at room temperature before addition of primary antibody and overnight incubation rocking at 4°C. The next day, four 500-750 microliter rinses with blocking solution, after pelleting with the mini microcentrifuge, preceded the addition of secondary antibodies in the dark, for two hours, at room temperature. Three short rinses with blocking solution after pelleting then followed before mounting 5 to 10 ovary pairs. The ovaries were teased apart on glass slides with tungsten needles or insect pins, and excess buffer was carefully removed by wicking with a tissue before addition of room temperature Prolong Gold Antifade Reagent with DAPI (Invitrogen) and No 1.5 glass coverslips.

Full-length *w*Mel WalE1 and *w*Ana FtsZ protein-encoding constructs were synthesized by GenScript using codons optimized for *E. coli* and cloned into pCR-TOPO before subcloning into the pUC57 expression vector (using BamHI and XhoI restriction enzymes). Sequence-verified constructs were expressed in *E. coli* BL21* and purified using Ni-NTA columns (Sigma-Aldrich). Because pUC57 contains a TEV cleavage site, N-terminal His-tags was removed from these proteins before sending them to Cocalico Biologicals for antibody generation. Rabbit polyclonal antibody sera against both full-length purified proteins were generated (Cocalico Biologicals, Inc) and used separately for immunohistochemistry (1:500 and 1:150, respectively). Secondary antibodies to rabbit and mouse IgGs that were highly purified to reduce cross reactivity were used with 488, 594 and 647 AlexaFluor conjugates at 1:1000 dilution in PBST with 0.5% BSA (Invitrogen ThermoFisher A11070, A-21244, A-21203, A-31571). For F-actin detection, we used Rhodamine-labelled Phalloidin or Acti-stain 488 Fluorescent Phalloidin (Cytoskeleton, Inc), per manufacturer instructions, depending on the cross and the wavelengths of fluorophores utilized.

Images were taken as Z-series stacks at 0.3 to 1.0 um intervals using a Nikon Ti2 fluorescent microscope with 60x oil objective and processed using NIS Elements software (Nikon). Care was taken such that exposure times were normalized across all experimental conditions. For quantification of *Wolbachia* within the developing oocyte, maximum projections stacks generated were used, excluding the peritoneal sheath. The irregular blob tool was used to outline entire egg chambers, using cortical actin staining as a guide, and DAPI DNA staining along with Actin staining was used to determine egg chamber stages.

### Co-Immunoprecipitation and Mass Spectrometry

Confluent *w*Mel-infected JW18 cells and uninfected TET-JW18 cells were harvested from 25 cm^2^, 50 mL non vented tissue-culture culture flasks. Cells were pelleted and washed three times with 1 ml volumes of PBS to remove culture medium and each cell type pellet was resuspended in 200 ul Lysis Buffer II (10mM Tris pH 7.4, 150mM NaCl, 10mM NaH2PO4, 1% Triton X-100) with added 1X Halt™ Protease Inhibitor Cocktail (ThermoFisher Scientific) and 5 mM EDTA on ice for 10 minutes, vortexing every 2 minutes. Debris and nuclei were pelleted at 10K rpm for 10 minutes in a microfuge (9391 g), 4°C rotor. Five microliters of rabbit pAb-WalE1 antisera were added to 100 microliters each of JW18 and JW18-TET lysate supernatants. A protein-A magnetic bead and column kit was used for co-immunoprecipitation and enrichment per manufacturer protocols with the following specific parameters: 30 minute 4°C incubation of immune complexes and magnetic protein A beads; prewetting columns with 200 microliters of 70% ethanol; and use of a stringent high-salt wash buffer (500 mM NaCl, 1% Igepal CA630 (NP-40), 50 mM Tris HCl (pH 8.0) before elution of proteins from the columns (Miltenyi Biotech, Inc). Eluted fractions were run 1-2 centimeters into a 4-20% PAGE minigel (NuSep) and the wedge of gel cut out from dye front to wells was submitted for analysis to the Indiana University Laboratory for Biological Mass Spectrometry. Peptides from *Drosophila melanogaster* and *Wolbachia pipientis* were identified and analyzed by LC-MS on an Orbitrap Fusion Lumos Tribrid equipped with an Easy NanoLC 1200.

### *Transgenic Drosophila stocks*, Wolbachia *infection detection and clearing*

All genetic backgrounds and fly stocks used in this study, including those obtained from the Bloomington Drosophila Stock Center are indicated in Supplementary Table 1. The *w*Mel infection status for all strains was determined by PCR of a *wsp* gene fragment or western blot of WSP protein (see Supplementary Table 1). Three Crispr guide RNAs were designed using the target finder at FlyCrispr (https://flycrispr.org/target-finder/) to the 5’ half of the Past1 gene for interruption of function in both the ubiquitous (B) and testes specific (A) splice variants. The three targets synthesized within the Crispr forward DNA primers were: Target 1 <u>3R:12698125..12698147</u> GAAGCGCG|AGAAGAACACCC; Target 4 <u>3R:12698221..12698243</u> GTTCCACGACTT|TCACTCGC and Target 6 <u>3R:12698304..12698326</u> CGGGCAAGACGA|CCTTCATC (Integrated DNA Technologies, Inc for forward and reverse primer synthesis). The guide RNAs were made per methods described in (44) and shipped out on dry ice to be mixed and injected into *Drosophila* embryos (Rainbow Transgenics, Inc). Two independent lines including a Past1 abrogation were recovered (Figure S1). Past1 primers used for amplification and sequencing through an 833 bp region that includes the three Crispr targets were: forward primer ATAACTGCCGTAGTCGTCGC (3R:12697988 - 12698007) and reverse primer GGAGCCGATGTAGACACGAG (3R:12698851 - 12698870). Standard methods were used for all crosses and culturing, and flies were maintained and crosses were performed at 22.5°C. Eleven stocks used in this study were obtained from the Bloomington Drosophila Stock Center (BDSC) at Indiana University (http://flystocks.bio.indiana.edu/), see Supplementary Table 1. Stocks carrying P{w[+mC]=UASp-RFP.WalE1} on the 2 or 3rd chromosomes (created in a *w*[1118] background and previously published (8)) were also used. Prior to use, the *Wolbachia* infection status of all stocks was investigated by pcr amplification of a ∼500 bp segment of the wMel *Wolbachia* surface protein *(wsp)* gene (wspF1 GTCCAATARSTGATGARGAAAC and wspR1 CYGCACCAAYAGYRCTRTAAA) and/or western blotting to detect the WSP protein. To make genetically similar stocks without *Wolbachia* infection, flies were treated with tetracycline and repopulated with other microbiome members as previously reported (45).

### Drosophila silver nitrate toxicity assays

Sets of 10-20 virgin females (uninfected, *w*[+] eyes) and 15-30 male flies were mated in small cages on standard fly food for two days and then allowed to lay eggs for 24 hours on grape agar (Genesee Scientific) plates (60 × 15 mm) with a little pile of yeast paste (1 g live baker’s yeast per 2 ml water) for 4-5 days. The 24 hour old plates were incubated at room temperature in a moist chamber another day until 48 hours, at which point 1st instar larvae were collected in a 100 micron disposable sieve, rinsed with room temperature water to remove food and debris, and were placed in groups of 10 larvae per plastic vial of 5 milliliters of minimal medium (5 g agar, 5 g dextrose, 360 milliliter distilled water, solubilized using heat and aliquoted) with ∼60 microliters of yeast paste made with water or a 0.003 % silver nitrate solution (Ag_2_NO_3_) applied to the surface of the minimal medium (based on method described in (32)). Vials were observed daily through the remainder of development. Pupal formation, adult emergence, and adult marker phenotypes were recorded for each vial through ∼19 days.

### Statistical analyses

All statistical analyses were performed in RStudio (v2022.07.01). Intensity data from Nikon NiE Elements were tested for normalcy and overdispersion in R. For yeast endocytosis assays, a Kruskal Wallis test was used. For *Wolbachia* titer in whole flies, quantified by western blot, normalized band intensities were used and an ANOVA applied to test for differences based on genotype. For *Wolbachia* titer in *Past1* mutant egg chambers, a GLM with a Quasipoisson distribution (glm in the stats library) was used to test for significant differences in the intensity of *Wolbachia* staining in each egg chamber across genotypes.

## Acknowledgements

This work was supported by a National Institutes of Health, National Institutes of Allergy and Infectious Diseases (NIAID) grant R01AI144430 to ILGN. We thank the Indiana University Laboratory for Biological Mass Spectrometry for their help in identifying peptides from our experiments.

## Supplementary Figures

**Figure S1.**
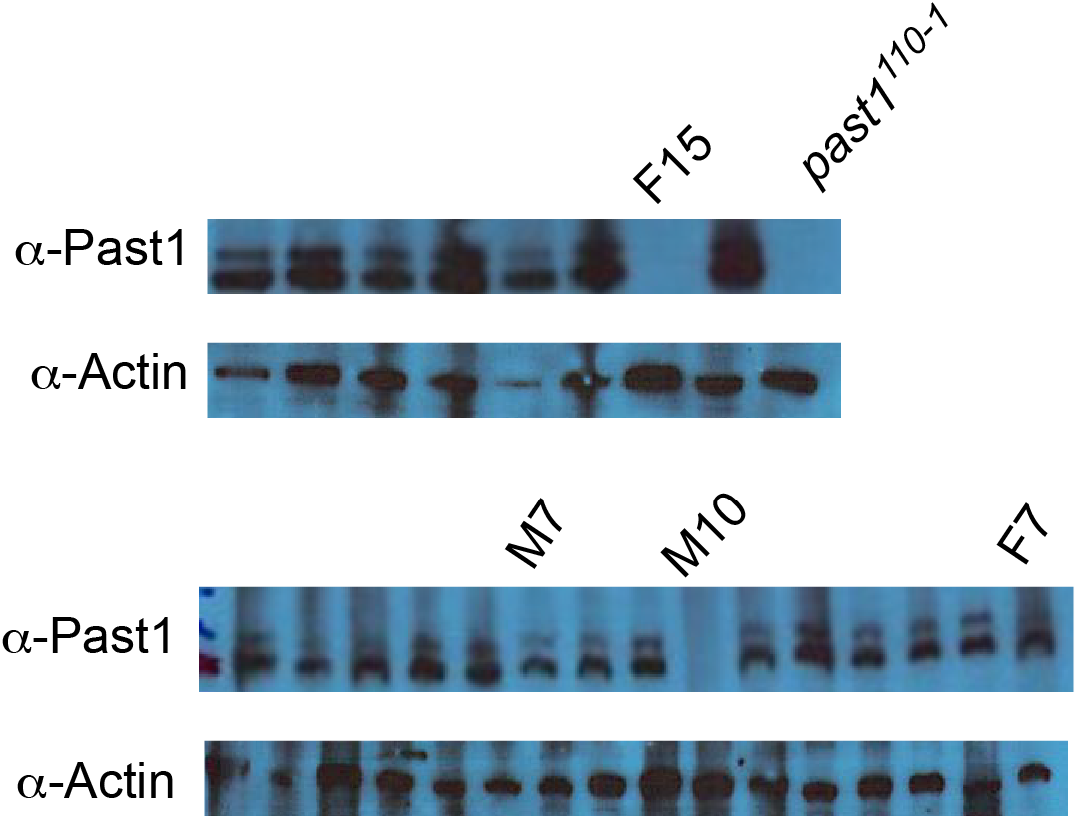
Past1 can no longer be detected by western blot in CRISPRKnock out flies M10 and F15. Past1^110-1^, M7, and F7 included as positive and negative controls, respectively (along with protein from other unmarked sibling fly lines that did not become null mutants in the Past1 gene following Crispr targeting).

